# The Susceptibility of Airborne SARS-CoV-2 to Far-UVC Irradiation

**DOI:** 10.1101/2025.06.09.658611

**Authors:** Darryl M. Angel, Irvan Luhung, Keyla S. G. de Sá, Jordan Peccia

## Abstract

Far-ultraviolet-C (far-UVC) irradiation has emerged as a breakthrough disinfection technology for the treatment of indoor air. Far-UVC wavelengths (222 nm) from filtered krypton-chloride excimer lamps are effective at inactivating airborne viruses and safe for human exposure, thus enabling the continuous treatment of bulk air in occupied settings. This study quantifies the susceptibility of airborne SARS-CoV-2, aerosolized in human saliva, to far-UVC radiation. We measured fluence rate-based Z value susceptibility constants (± std. err.) of 4.4 ± 0.6 and 6.8 ± 0.7 cm^2^ mJ^-1^ for airborne SARS-CoV-2 under 40% and 65% relative humidity (RH) levels, respectively. At modeled far-UVC irradiation levels corresponding to 25% of the maximum safe human exposure limit, the resulting far-UVC equivalent air changes per hour (eACH) exceeded 62 hr ¹ at 65% RH and were significantly greater than the corresponding airborne SARS-CoV-2 natural decay rate (( std. err.) of 5.4 ± 1.1 hr ¹, measured in the absence of far-UVC. These results define first-order loss rates for airborne SARS-CoV-2 under far-UVC exposure and support quantitative risk assessments and rational disinfection system implementation.

**Synopsis Statement:** Quantified the ability of far-UVC to facilitate airborne SARS-CoV-2 inactivation, guiding the implementation of far-UVC to promote transmission risk reduction in occupied spaces.

## 1. INTRODUCTION

Severe Acute Respiratory Syndrome Coronavirus 2 (SARS-CoV-2) infections have resulted in negative economic, educational, and social impacts, were responsible for nearly 7 million deaths worldwide from 2020 to the end of 2023, and is now endemic with new strains continuing to emerge.(1–3) As the importance of airborne transmission for SARS-CoV-2 has been clarified, effective environmental intervention and engineering controls hold renewed potential for interrupting virus exposure and mitigating subsequent outbreaks of SARS-CoV-2 and other airborne viruses.(3–7) While common environmental approaches to reduce airborne transmission of SARS-CoV-2, including ventilation, portable air filtration, and upper-room ultraviolet germicidal irradiation (UGVI), have demonstrated effectiveness towards reducing respiratory virus concentrations in indoor air, their implementation has been hampered by cost, efficiency, or their limited applicability in congregate settings.(7–20)

Treatment of bulk air by far-ultraviolet irradiation (far-UVC) has emerged as a safe technology to effectively inactivate airborne pathogens in occupied spaces, circumventing many of the above limitations.(15, 21–24) Far-UVC irradiance is commonly produced by krypton chloride (KrCl) excimer lamps, primarily emitting 222 nm wavelength light.(23, 24) This wavelength cannot penetrate the outer protective layers of human skin and eyes tissue, enabling the utilization of far-UVC for the treatment of bulk-air within occupied settings.(18, 23) Current literature evaluating the effectiveness of UV irradiation reports similar or improved inactivation of airborne viruses, including human coronavirus (HCoV) OC43, when subjected to 222 nm compared to conventional 254 nm UVGI.(18, 22, 25, 26) Further, across comparisons of 222 nm through 365 nm UV light generating systems, the far-UVC KrCl lamp required the lowest electrical energy consumption to facilitate 1 log inactivation of aerosolized viruses.(18)

Experimental parameters, including relative humidity (RH) and aerosolization liquid, have been identified as significant factors that influence the performance of UV-mediated disinfection on viral aerosols.(26–28) The high energy wavelength of far-UVC technology may promote the formation of reactive oxygen species (ROS) in more hydrated particles, indicating the potential for increased inactivation rates at elevated RH levels.(29) However, the relationship between 222 nm airborne viral inactivation and RH remains uncharacterized.

The installation of far-UVC devices spread during and following the COVID-19 pandemic, yet, while far-UVC driven inactivation of SARS-CoV-2 in dried droplets has been explored (30), no experimental studies exist that evaluate the susceptibility of airborne SARS-CoV-2 to far-UVC irradiation. Susceptibility constants are needed for the rationale design of far-UVC implementation and to evaluate the associated transmission risk reduction. This study quantifies the susceptibility of airborne SARS-CoV-2 to 222 nm far-UVC via chamber aerosolization experimentation. Experiments were conducted in a Biosafety Level 3 (BSL-3) laboratory, exposing airborne SARS-CoV-2 to varying doses of 222 nm irradiation under ∼40% and ∼65% relative humidity levels. The susceptibility of SARS-CoV-2, aerosolized in human saliva, was evaluated by reporting the far-UVC driven inactivation rate constants, *Z* value susceptibility constants, and natural decay rate constants. Potential infection risk reduction in congregate settings was modeled as a function of far-UVC dose and building RH.

## 2. MATERIALS AND METHODS

### 2.1 SARS-CoV-2 Propagation and Quantification

Details regarding the propagation and quantification of infectious and total SARS-CoV-2 beta variant (B.1.351) are explained in the Supplementary Information (SI). Briefly, SARS-CoV-2 was propagated on Vero E6-TMPRSS2-T2A-ACE2 cells (#NR-54970, BEI Resources) that were cultured in Dulbecco’s Modified Eagle Medium (DMEM) (#11965092, Gibco) supplemented with 10% heat-inactivated fetal bovine serum and 1mM sodium pyruvate at 37°C in a 5% CO_2_ incubator.(31, 32) Puromycin (0.01 mg mL^-1^, final concentration) was added to the cell culture DMEM to select for ACE2 and TMPRSS2 expression in Vero E6 cells.(33) Post sampling, collected SARS-CoV-2 aerosols were quantified using droplet digital PCR (ddPCR) and plaque assay (PA) methods, as previously described.(31, 34) The ddPCR-quantified SARS-CoV-2 represent the total viral aerosols collected, enumerating both infectious and inactive viral aerosols. In contrast, SARS-CoV-2 concentration enumerated by plaque assay –on Vero E6 cells in 6-well plates– are representative of solely the still-infectious virus collected.(31) As such, ddPCR-based decay, or total viral decay, is inclusive of depositional mechanisms and potential air leakage, while plaque assay-based decay (PA), or infectious decay, is attributable to losses of infectious viral aerosols resulting from inactivation in addition to the aforementioned mechanisms.

### 2.2 Experimental Setup

#### 2.2.1 Aerosolization Chamber

A sterilized 64 L cubic chamber was utilized for aerosolization experiments (Figure 1) inside a Class II Biosafety Cabinet in a BSL-3 laboratory. The aerosolization chamber housed a far-UVC KrCl excimer light source (Care222 B1.5 Illuminator, Ushio America, Inc.) within a 4 L bump-out in chamber, separated from the viral aerosols via 1/8" thick quartz plate, allowing for sufficient UV transmission.(35) The B1.5 far-UVC device module, containing a filter to block light wavelengths greater than 230 nm, was utilized as it contains a diffuser to facilitate a 110 emission angle to more effectively distribute far-UVC irradiance throughout the experimental chamber. One 240 µm UV semi-transparent plastic sheet (EUROPLEX® Film 0F304, TOPAS 8007 resin, Polyvantis Sanford LLC) was secured in front of the far-UVC device, separated by the quartz plate, to reduce the far-UVC dose received by the viral aerosols (4.1% transmittance) and allow for observed viral decay across multiple time points without altering the light distribution.(21, 36) The main chamber housed two fans to promote well mixed conditions and a relative humidity and temperature probe (HOBO).(37) The fans had 4”-sized blades that produced an air speed of ∼4 m sec^-1^. Based on our chamber’s volume and the volumetric air movement provided by the two fans placed on opposite sides of the chamber, a well-mixed condition was expected to be reached within seconds.

**Figure 1.**
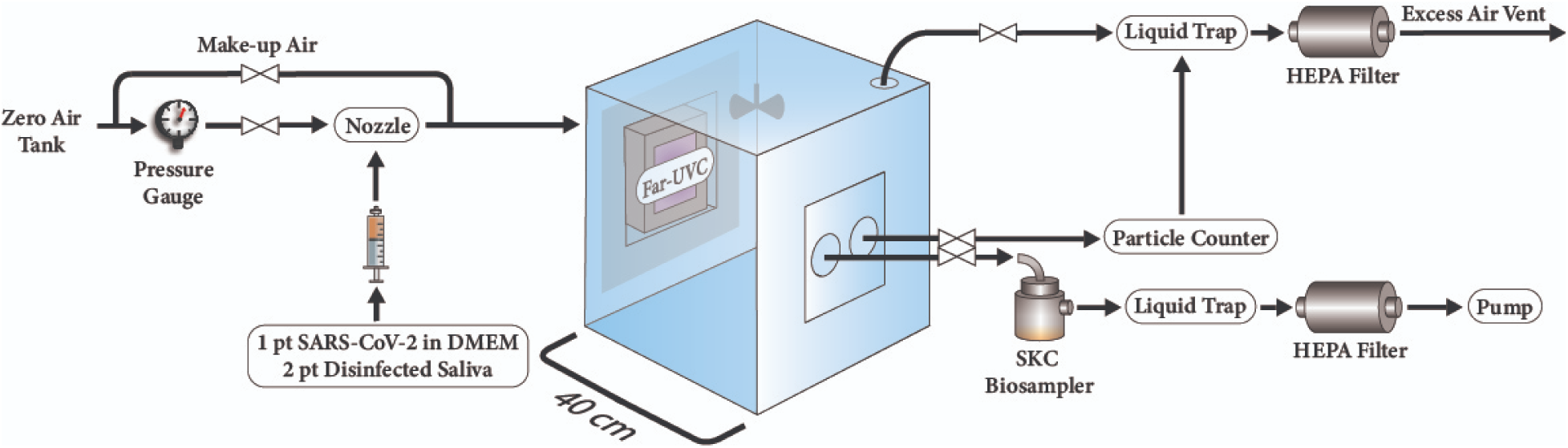
Aerosolization chamber housing KrCl excimer lamp far-UVC light source and sampling setup. The far-UVC device is housed within a bump-out, sectioned off from the chamber using a quartz glass plate. The open circles in the plexiglass sampling block represent sampling ports fitted with on/off valves for the SKC BioSamplers and optical particle counter as well as an exhaust port fitted with a liquid trap and HEPA filtration.

#### 2.2.2 Experimental Procedure

SARS-CoV-2 viral aerosols (∼10^7^ PFU mL^-1^), in a 2 part UV-disinfected human saliva (#IRHUSL5ML, Innovative Research) and 1 part DMEM solution (rationale in SI), were generated via air assisted atomizing nozzle operating at 20 PSI (#IAZA500415K, The Lee Company) and aerosolized into the chamber in one batch at the experimental onset. Three viral aerosol sample time points were considered per experiment by collection into liquid impingers (#225-9595, BioSamplers, SKC Inc.) filled with 15 mL of sterile DMEM at a flow rate of 12.5 L min^-1^. A filtered make-up air line preventing vacuum conditions during collection. Total particle counts were taken in between viral bioaerosol time points by an optical particle counter (MET ONE HHPC 6+). For far-UVC on experiments, the excimer lamp was turned on to stabilize for at least 5 minutes prior to viral aerosolization.

Two experimental RH conditions, deemed low RH and high RH, were explored. Chamber relative humidity was set by the amount of 1:2 SARS-CoV-2 in DMEM and saliva aerosolized. Each experimental condition (far-UVC_on_ at low RH, and far-UVC_off_ at low RH, far-UVC_on_ at high RH, far-UVC_off_ at high RH) was conducted in independent duplicate experiments. The temperature in the chamber remained stable at a mean (± standard error) of 25.2 ± 0.1 .

#### 2.2.3 Measurement and Modeling of Chamber Irradiance and Fluence Rate

To model the irradiance distribution and estimate the fluence rate experienced by aerosols within the chamber, chamber dimensions (5% wall reflectance) (38) and the B1.5 Illuminator far-UVC light source were input into UV simulation software (OSLUV Illuminate).(39) This modeling software utilizes the default B1.5 Illuminator output, thus, scaling factors were required to account for the reduction in irradiance facilitated by the semi-transparent sheet. A UIT2400 Light Meter (Ushio America, Inc.) was used to measure the irradiance inside the chamber along the front, middle, and back planes, with the light meter facing the far-UVC lamp wall.(40) All irradiance measurements were taken at room temperature and ∼50% RH. Direct light meter measurements at the center point of each of the three vertical planes were then used to produce dimensionless scaling factors which were applied to obtain modeled chamber irradiance at a higher resolution than practically measurable, as well as the average irradiance and fluence rate within the chamber (see SI).

To estimate the fluence rate within a realistic congregate setting, a 30 x 20 x 10 ft classroom, the OSLUV Illuminate model was used to determine the average fluence rate corresponding to 25% of the American Conference of Governmental Industrial Hygienists (ACGIH) threshold limit value (TLV) for safe human eye exposure to far-UVC irradiation. In the model, Ushio B1.5 KrCl excimer lamps were evenly distributed onto the ceiling of the simulated classroom, inclusive of floor, celling, and wall paint reflectance (7.0%, 7.1%, 8.1%, respectively).(11, 39) Within this classroom scenario, the simulated installation of 26 USHIO B1.5 units on the ceiling in an equally distributed manner yields ∼25% of the current TLV; as such, it is impractical to report SARS-CoV-2 transmission reduction associated with exposure to far-UVC at the TLV of 161 mJ cm^-2^ in the classroom scenario.

### 2.3 Viral Aerosol Disinfection

#### 2.3.1 Far-UVC Mediated Inactivation Decay Rate Constant and Susceptibility Constant

First-order decay rate constants were obtained as the slope of a least-squares linear regression line on a plot of the natural logarithm of the viral concentration as a function of time.(37, 41, 42) Following normalization based on initial SARS-CoV-2 concentration, to account for the varying concentrations aerosolized, the pooled viral decay from independent duplicate experiments was used to define each rate constant. The inactivation decay rate constant of SARS-CoV-2 solely attributable to far-UVC exposure, *k_inactivation_*, can be calculated using the experimentally derived infectious and total (RNA-based) decay rates of viral aerosols under far-UVC on conditions, defined as *k_PA_* and *k_ddPCR_*, respectively (Equation 1). Subtracting *k_ddPCR_* and the background natural infectious decay, *k_natural decay_*, from the infectious decay rate of far-UV on experiments ensures the removal of depositional, potential air leakage, and natural infectious viral decay factors.(40)

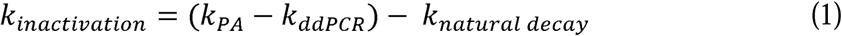

Requiring both plaque assay and ddPCR viral aerosol quantification, the natural infectious decay rate of SARS-CoV-2 within the experimental setup, *k_natural decay_*, is defined as the infectious viral decay rate constant minus the total viral decay rate constant under background (far-UVC off) conditions. The far-UVC mediated viral inactivation rate constant was calculated for both of the experimental relative humidity conditions, with each decay rate constant in the units of hr^-1^, or equivalent air changes per hour (eACH). Utilizing the determined eACH, the clean air delivery rate (CADR), which is traditionally reported as ft^3^ min^-1^ (CFM), is calculated as the product of the technology-mediated decay rate constant, or eACH, and the chamber volume.(43)

To enable the application of quantified SARS-CoV-2 aerosol disinfection relative to far-UVC exposure, the UV susceptibility constant, or *Z* value (cm^2^ mJ^-1^), was calculated using the average fluence rate or average irradiance (*F* or *I*, mW cm^2^) and the inactivation rate constant in the units of sec^-1^ (Eq. 2).(18, 40, 44)

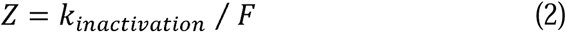

While the fluence rate, or spherical irradiance, is relevant to the inactivation of viral aerosols, susceptibility constants derived from irradiation are commonly reported in published literature. Thus, irradiance-based *Z* values are reported to enable comparisons with existing literature, while all risk of infection and airflow scaling calculations were conducted utilizing the fluence rate associated *Z* values.

#### 2.3.2 Risk of Infection

The risk of infection, or transmission risk, was estimated using an open source adjusted Wells-Riley model (v. 3.6.8), calibrated based on COVID-19 literature for quanta emission rate.(45–48) This risk of infection represents the probability, as a percentage, that one individual will be infected with SARS-CoV-2 from a singular event within the envisioned scenario (48); a 30 x 20 x 10 ft classroom occupied by 29 susceptible people and 1 infected person, with no individuals wearing a mask (one-time event, 1 hour). The default quanta exhalation rate was scaled to account for the SARS-CoV-2 variant (3.3-fold, Omicron BA.2) and respiratory activities (4.7-fold, speaking). Within the modeled scenario, a ventilation rate of 3 hr^-1^ was utilized, while the natural decay rate and deposition rate of airborne SARS-CoV-2 were obtained from the experimental results presented in this study and scaled to the size of the classroom. The additional disinfection rate (hr^-1^) of airborne SARS-CoV-2 provided by far-UVC exposure was estimated from the fluence-based susceptibility constants (*Z* values) derived from our experiments for low and high RH conditions. The event reproduction numbers (R_e_), representing the average number of new infections that can be caused by a single individual, were estimated for this scenario from an existing model by Tupper et al.(49).When R_e_ < 1, the outbreak is receding.

### 2.4 Statistics

Least squares linear regression, constraining the y-intercept to zero, was conducted to fit experimental data to a first-order decay rate model. To assess the statistical difference within and across experimental RH conditions, a Kruskal Wallis test followed by Dunn’s Multiple Comparison test was utilized. Analysis was conducted in GraphPad Prism v. 10.1.2 using two-tailed methods.

## 3. RESULTS

### 3.1 UV Irradiance and Fluence Rate

Based on UV light simulation, the average far-UVC fluence rate within the chamber was estimated to be 0.96 µW cm^-2^ while the average far-UVC irradiance was 0.70 µW cm^-2^. As depicted in Figure 2, there was notable spatial heterogeneity of irradiance in the chamber within and across the three vertical planes modeled, ranging from 0.07 to 7.69 µW cm^-2^. This gradient of 222 nm irradiation also exists in any current application of far-UVC technology. However, under the well mixed conditions within our chamber the far-UVC irradiation experienced by SARS-CoV-2 aerosols is assumed to be equivalent to the average values above.

**Figure 2.**
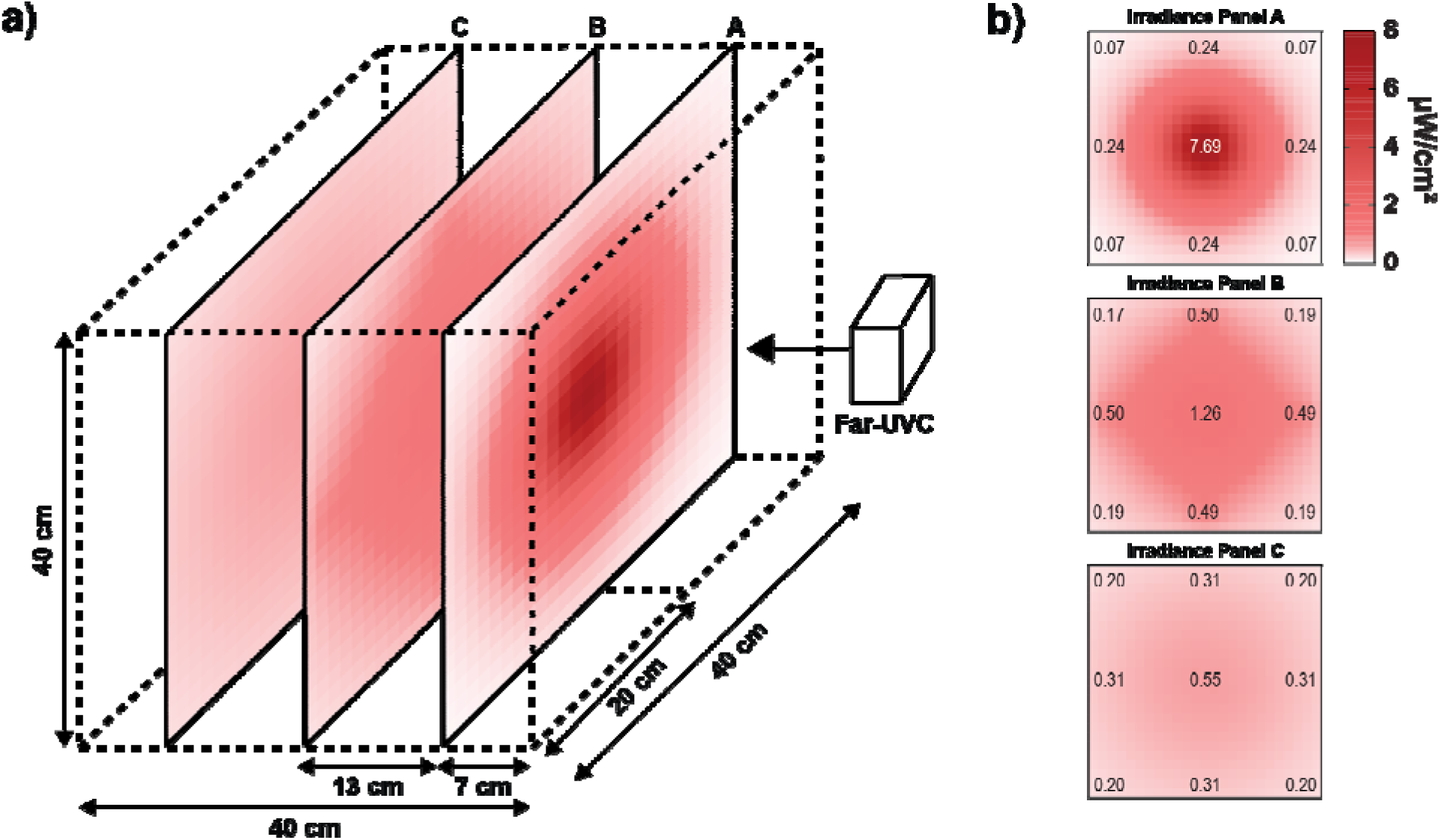
Modeled spatial distribution of 222 nm irradiance within the chamber. a) Spatial distribution of irradiance within the cubic aerosolization chamber, with the far-UVC lamp on the right edge of the chamber boundary. b) Irradiance panels A, B, and C demonstrate a front-facing perspective of the vertical 222 nm irradiance within the chamber along the closest (7 cm), middle (21 cm), and furthest (33 cm) vertical planes from the far-lamp, respectively. The center points of each panel represent the measured irradiance values.

### 3.2 SARS-CoV-2 Susceptibility to Far-UVC

The average experimental relative humidities for low and high RH conditions (± standard error) are 38.6 ± 0.4% and 67.1 ± 0.6%, respectively (Table S1). The geometric mean diameter (± geometric standard deviation) of chamber aerosols produced by the atomizer is 1.38 ± 2.06 µm, remaining consistent across each experimental condition. This distribution (Figure S1) of aerosols produced, in the low micron range, is within the size range of human-exhaled respiratory viral particle emissions.(8, 50–52)

#### 3.2.1 Experimental Total and Infectious Viral Decay

Following chamber experimentation, first-order total and infectious SARS-CoV-2 aerosol decay rate constants under far-UVC off (background) and experimental far-UVC on conditions for both low and high relative humidities were determined (Figure 3). As shown in Figure 3, the normalized decay rate of infectious viral aerosols increases from far-UVC off to far-UVC on conditions (*p* ≤ 0.01) while remaining statistically insignificant for total viral aerosol decay (*p* = 0.32 under low RH; *p* = 0.28 under high RH), indicating the ability of far-UVC irradiance to facilitate inactivation of airborne SARS-CoV-2.

**Figure 3.**
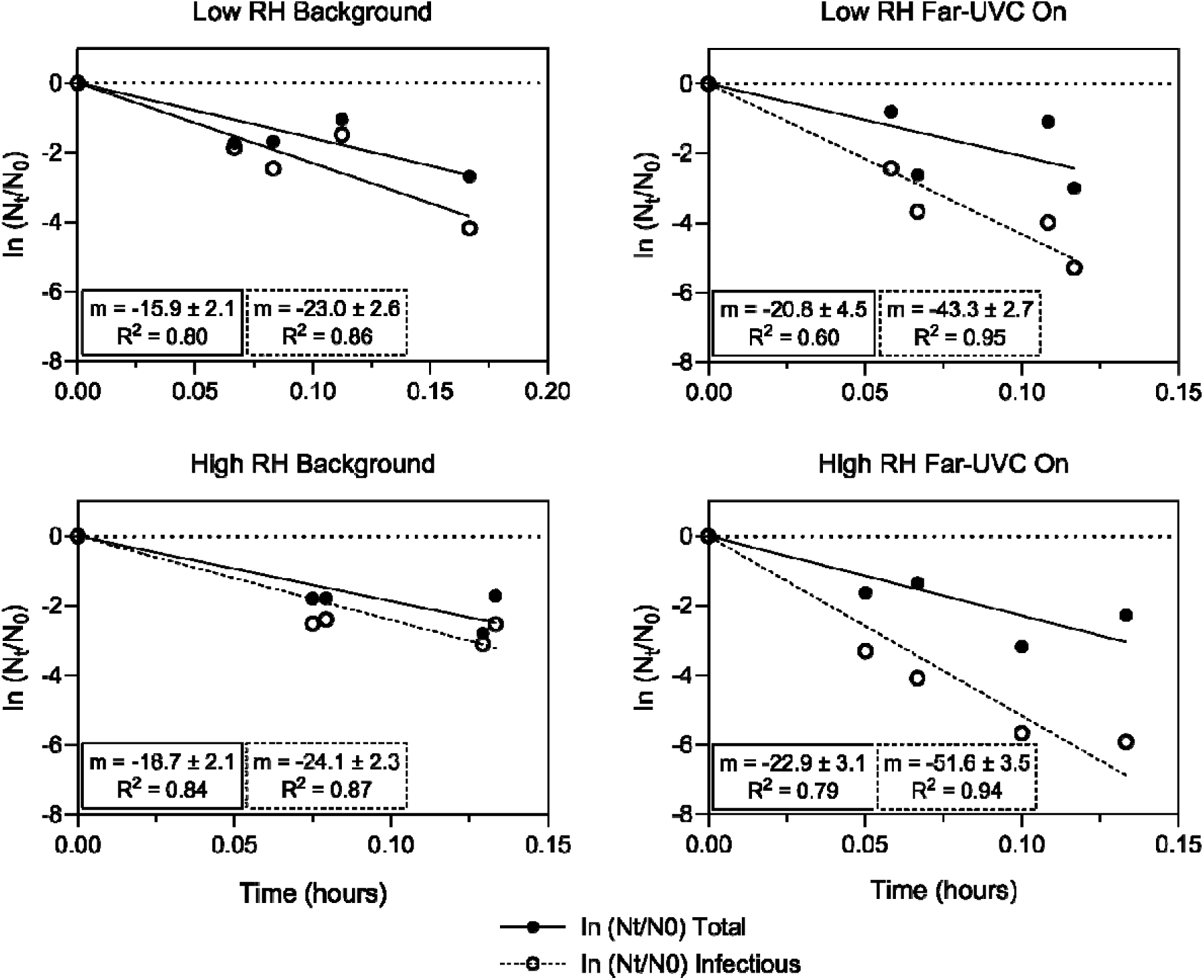
Linear regression analysis on total and infectious aerosolized SARS-CoV-2 exposed to 222 nm irradiation. Regression analysis on the natural logarithm of the normalized total (black circles) and infectious (open circles) SARS-CoV-2 concentration decay as a function of time from independent duplicate experiments. Slopes (m = slope) of these linear regression lines (± std. err.) identify the first-order decay rate constants (hr^-1^) for total and infectious SARS-CoV-2.

#### 3.2.2 Far-UVC-Driven Inactivation Decay Rates and Susceptibility Constants

Within the two experimental RH conditions, the total and infectious decay rate constants from far-UVC off and far-UVC on conditions can be used to calculate the viral aerosol inactivation rate constants attributable to elevated doses of far-UVC, as described in Equation 1. Accounting for the removal of airborne SARS-CoV-2 from natural infectious decay and depositional mechanisms, the decay rate constants solely attributable to far-UVC induced inactivation of SARS-CoV-2 (± standard error) are 15.3 ± 1.9 hr^−1^ under low RH and 23.4 ± 2.3 hr^−1^ under high RH, following average fluence rate exposure of 0.96 µW cm^-2^. A ∼2.1 and ∼4.3-fold increase was observed in the far-UVC driven SARS-CoV-2 inactivation over the natural infectious decay for low (*p* ≤ 0.01) and high (*p* ≤ 0.01) relative humidities, respectively (Figure 4). Additionally, there is an increase in far-UVC mediated viral inactivation rate constants associated with far-UVC exposure from low RH to high RH (*p* < 0.05).

**Figure 4.**
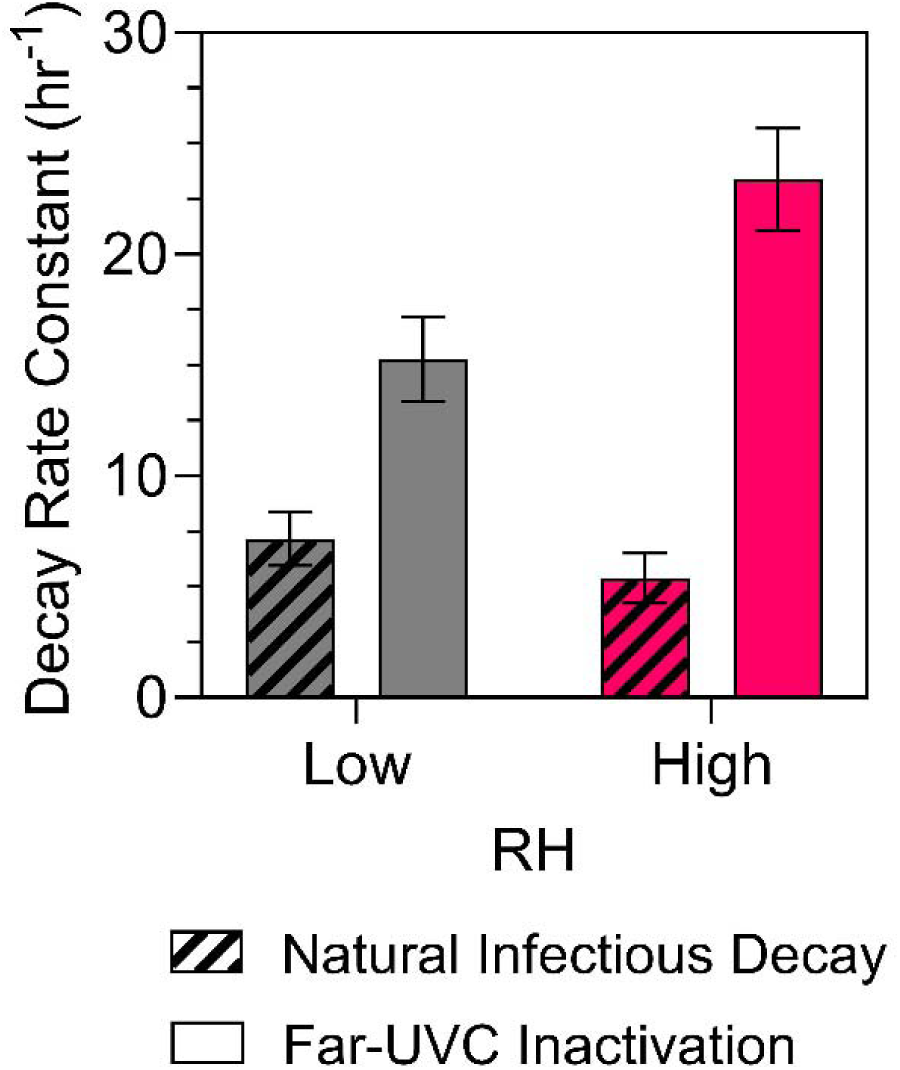
Natural infectious decay and far-UVC driven inactivation decay rate constants (± SE) for aerosolized SARS-CoV-2. First-order decay rate constants for the natural infectious decay (slanted bars) and far-UVC induced inactivation (solid bars) of SARS-CoV-2 aerosols under low (grey) and high (red) relative humidity conditions. The average 222 nm fluence rate was 0.96 µW cm^-2^ during far-UVC on experiments.

The average fluence rate and far-UVC inactivation decay rate constants result in airborne SARS-CoV-2 far-UVC susceptibility constants (± std. err.) of 4.4 ± 0.6 and 6.8 ± 0.7 cm^2^ mJ^-1^, for low and high RH, respectively. Table 1 demonstrates a comparison of viral susceptibility constants to far-UVC irradiation. Rather than the above *Z* value calculation utilizing fluence rate, which represents a more accurate characterization of far-UVC mediated inactivation of aerosols, far-UVC susceptibility constants for this study presented in Table 1 are derived from the average chamber irradiance, yielding *Z* values of 6.1 ± 0.8 (std. err). and 9.3 ± 0.9 cm^2^ mJ^-1^, only to provide comparisons with existing literature. The dose required to facilitate a 2-log decay in infectious SARS-CoV-2 aerosols, derived from fluence rate-based *Z* values, is 1.04 mJ cm^-2^ under low RH and 0.68 mJ cm^-2^ under high RH conditions. The 2-log decay irradiance-based doses are presented in the SI.

**Table 1.**
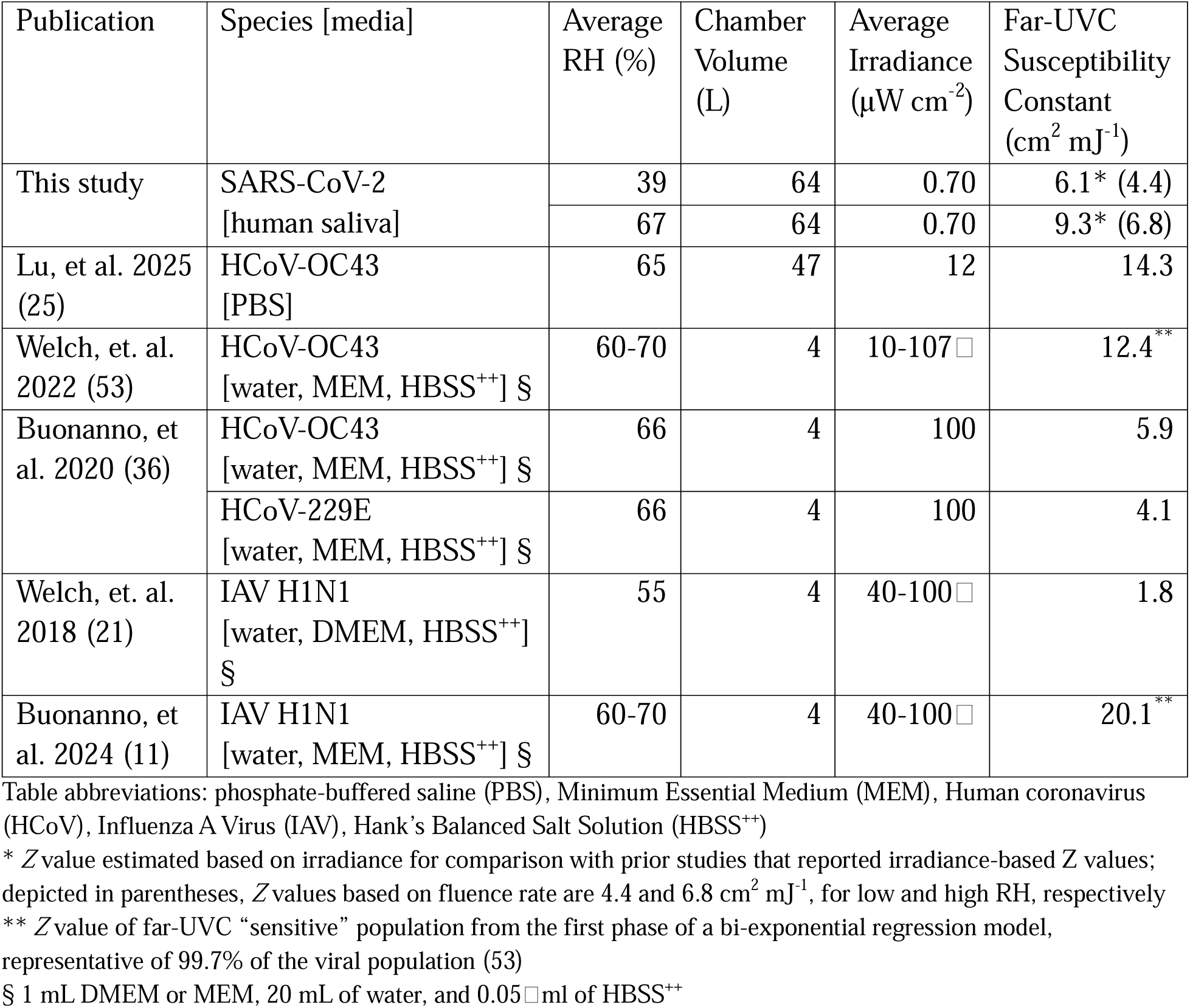
Experimental conditions and irradiance-based far-UVC susceptibility of aerosolized enveloped human viruses.

#### 3.2.3 Far-UVC Mediated SARS-CoV-2 Disinfection

To quantify 222 nm facilitated inactivation of airborne SARS-CoV-2 beyond the test chamber used in this study, the CADR can provide a metric for comparing the effectiveness of air cleaners as a function of room size.(43) Scaling for a 30 ft x 20 ft x 10 ft classroom setting, the Environmental Protection Agency (EPA) recommends a target minimum CADR of 390 cubic feet per minute (CFM) for air cleaners in a 600 ft^2^ room (54); reaching this guideline with far-UVC technology would require average fluence rates of approximately 0.25 µW cm^-2^ and 0.16 µW cm^-2^ for low and high RH conditions, respectively. This theoretical target mean fluence rate within the classroom scenario is well within the range of room fluence rates that are currently plausible, as a mean fluence rate of 0.82 µW cm^-2^ has been previously reported in a room setting employing far-UVC KrCl lamps.(11)

An adjusted Wells-Riley model was employed to further evaluate the effective implementation of far-UVC in occupied spaces relative to increasing 222 nm average fluence rates (Figure 5). Based on the SARS-CoV-2 adjusted Wells-Riley model using the parameters and scenario described in section 2.3.2, the baseline risk of infection (without far-UVC) associated with SARS-CoV-2 transmission from one emitter in a well-mixed classroom with 3 ACH from ventilation is 6.7% under low RH and 7.6% under high RH conditions, with the event reproduction number exceeding one. The event reproduction number for this modeled classroom scenario falls below one when the risk of SARS-CoV-2 infection is less than 3.5%, occurring at average fluence rates of 0.76 and 0.57 μW cm^-2^ under low and high RH, respectively.

**Figure 5.**
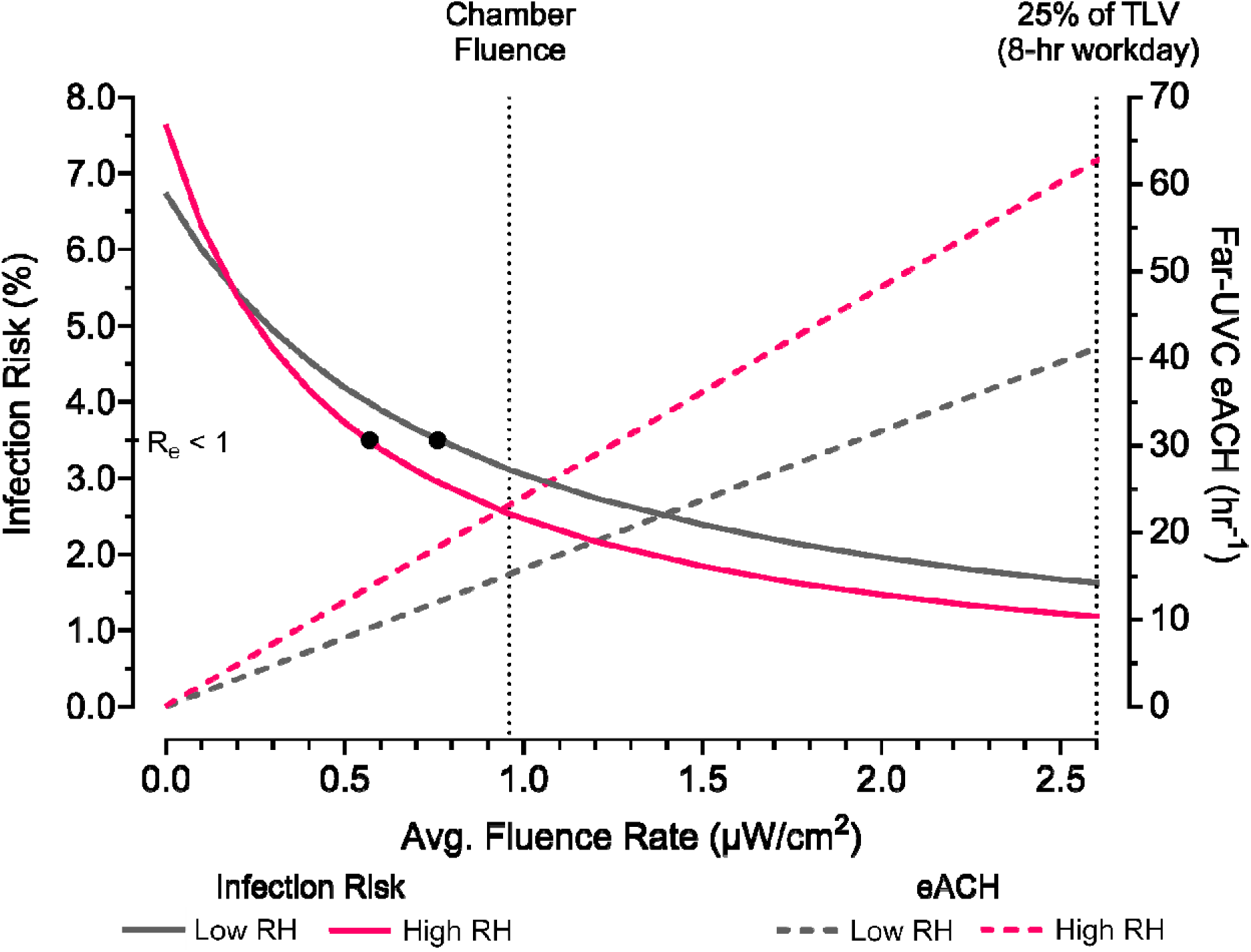
Probability of SARS-CoV-2 infection within the classroom scenario and eACH as a function of the far-UVC fluence rate across experimental RH conditions. The left y-axis depicts the risk of infection provided in a classroom setting, with 30 occupants and one emitter that is speaking, under a background ventilation level of 3 ACH, as a function of increasing average 222 nm fluence rate within the room. The right y-axis demonstrates the far-UVC driven eACH associated with 222 nm fluence rate. The black circles depict the risk of infection corresponding to the average fluence rate in which the event reproduction number (R_e_) is below one within the classroom scenario. Vertical markers demonstrate the average fluence rate utilized in this study and the average fluence rate corresponding to 25% of the TLV for safe human eye exposure within the simulated classroom.

When applying the previously reported attainable room-sized far-UVC average fluence rate of 0.82 μW cm^-2^ (11), this risk of infection within the classroom scenario decreases to 3.4% and 2.8% for low and high relative humidities, respectively. Based on the *Z* values obtained in the present study, the above fluence rate corresponds to far-UVC driven eACH/SARS-CoV-2 aerosol inactivation decay rate constants of 13.0 hr^-1^ under low RH and 19.8 hr^-1^ under high RH conditions. Using the defined classroom parameters, these eACH values translate to airflows of 43 and 66 CFM per person, which are higher than the 40 CFM per person minimum equivalent clean airflow recommended guideline from the ASHRAE Standard 241 for Control of Infectious Aerosols.(55)

It is also pertinent to assess the transmission risk relative to the threshold limit value for average far-UVC exposure in an occupied indoor space. Within the classroom volume, an average fluence rate of 2.60 μW cm^-2^ corresponds to approximately one quarter of the 161 mJ cm^-2^ TLV for safe human eye exposure to far-UVC irradiance (1.8 m height) in eight-hour work day.(18, 22–24, 40) At 25% of the maximum allowable 222 nm UVC exposure, the associated risk of SARS-CoV-2 infection is reduced to 1.62% and 1.18% under low and high RH conditions, respectively. The corresponding far-UVC mediated eACH increases to 41.2 hr^-1^ at 39% RH and 62.7 hr^-1^ at 67% RH.

## 4. DISCUSSION

Recent revisions to the ACGIH standard for exposure to far-UVC at 222 nm place threshold limit value at 161 mJ cm^-2^ for eye exposure and 479 mJ cm^-2^ for skin exposure in an eight hour occupational day.(22–24, 40) The TLV’s of 254 nm are at least 26-fold lower than 222 nm, outlining the decreased adverse human health effect risks of far-UVC when compared to conventional UVGI and enabling the safe implementation of far-UVC for pathogen disinfection in occupied spaces.(15, 22) Far-UVC technology is considered a safer alternative for in-room irradiation owing to the improved absorbance of 222 nm by proteins and subsequent limited penetration into the corneal epithelium and stratum corneum, the outermost layers of human eye and skin tissue.(15, 16, 18, 23)

Aerosols and droplets can experience 222 nm irradiation from all sides, thus, the *Z* values presented for SARS-CoV-2 inactivation by far-UVC quantified as a function of fluence rate, or spherical irradiance, provides an accurate measure for the susceptibility of such bioaerosols. The irradiance-based *Z* value susceptibility constants are derived from 222 nm irradiation experienced from one plane, better serving as a measure for disinfection among surface and potentially aqueous environments. As the average fluence rate estimated within the chamber is greater than the average irradiance, at 0.96 µW cm^-2^ and 0.70 µW cm^-2^, respectively, the fluence-based *Z* values (4.4 and 6.8 cm^2^ mJ^-1^) reported for SARS-CoV-2 inactivation under low and high RH conditions are less than those reported for the irradiance-based *Z* values (6.1 and 9.3 cm^2^ mJ^-1^), given that the UV susceptibility constant is calculated as the quotient of the inactivation rate constant and average fluence-rate or irradiance (Eq. 2).(18, 40, 44)

The airborne SARS-CoV-2 irradiance-based susceptibility constants of 6.1 and 9.3 cm^2^ mJ^-1^ for low and high relative humidities, respectively, are within the range of far-UVC viral aerosol susceptibility constants reported in the current literature.(11, 18, 19, 21, 22, 24, 25, 36, 53). At relative humidities of 60-70%, comparable studies report far-UVC susceptibility constants of 4.1–14.3 cm^2^ mJ^-1^ for HCoV-OC43 and HCoV-229E aerosolized in a supplemented water and infection medium solution or PBS.(25, 36, 53) The susceptibility constant of 9.3 ± 0.9 cm^2^ mJ^-1^ for SARS-CoV-2, aerosolized in saliva, obtained in the present study under an average RH of ∼67% is within this range of far-UVC susceptibility constants for human coronaviruses.(25, 36) Similarly, the *Z* values calculated here for SARS-CoV-2 are commensurate with those observed for far-UVC driven inactivation of the non-coronavirus organisms influenza A and Ф6.(11, 18, 21, 24, 26) Like SARS-CoV-2, both IAV and Ф6 are enveloped RNA viruses.(34, 56, 57) Compared to the viral surrogate MS2, which has been suggested as a model organism for airborne SARS-CoV-2 inactivation (58–61), the SARS-CoV-2 far-UVC *Z* values in the present study are greater than those reported for MS2 (18, 26), identifying the enhanced susceptibility of SARS-CoV-2 to far-UVC disinfection. This comparison suggests the important role of a viral lipid envelope in far-UVC susceptibility and broadly matches the demonstrated improved resistance of non-enveloped viruses to UVC inactivation.(25, 26, 62, 63)

Here, the observed far-UVC mediated airborne SARS-CoV-2 inactivation decay rate constants and susceptibility constants increase from low (39%) to high (67%) RH. This statistically significant difference in RH-dependent far-UVC driven SARS-CoV-2 decay provides a foundation to support further investigation under additional RH conditions to identify potential trends and clarify mechanistic changes. While additional comparisons among 222 nm disinfection of viral aerosols as a function of RH remain to be reported, this data indicates that relative humidity may be a significant factor in affecting far-UVC disinfection, as has been demonstrated for 254 nm UVGI.(27, 64) Past literature, however, does not present a unified theory regarding the impacts of RH on UV-mediated disinfection of viruses.(14, 27, 44, 64–68) Though RH appears to influence UV-254 airborne viral inactivation in a predicable manner for enveloped, single stranded viruses (14, 28, 65), the known mechanisms of inactivation by far-UVC differ from 254 nm photoinactivation, which predominately occurs via genome damage through the formation of pyrimidine dimers.(12, 13, 25, 69) In addition to genome damage, hypothesized to result from photodimerization, strand breaks, and/or the interruption of 3D structures of RNA, exposure to 222 nm irradiation further damages a variety of biomolecules and is thought to inactivate the virus through disruption of viral proteins and lipids.(25, 70) Beyond these mechanisms, past studies have indicated that the high energy of far-UVC has greater potential to excite and create free radicals, including ROS, with increasing particle hydration.(25, 29, 70) Subsequently, this may increase lipid peroxidation and further damage the lipid bilayer of the viral envelope, facilitating greater inactivation at higher RH (25, 70), though this specific far-UVC driven inactivation mechanism relative to RH remains to be evaluated.

This study quantifies the susceptibility of SARS-CoV-2, aerosolized in human saliva, to far-UVC (222 nm) irradiation. Far-UVC significantly enhanced viral aerosol inactivation over background conditions at both low (39%) and high (67%) relative humidities studied, coinciding with fluence rate-based far-UVC susceptibility constants of 4.4 ± 0.6 and 6.8 ± 0.7 cm^2^ mJ^-1^ and irradiance-based susceptibility constants of 6.1 ± 0.8 and 9.3 ± 0.9 cm^2^ mJ^-1^, respectively.

Exposure to far-UVC demonstrated improved 222 nm driven inactivation at high RH relative to low RH. Furthermore, far-UVC induced SARS-CoV-2 aerosol inactivation reaches equivalent air exchange rates of greater than or equal to 41.2 hr^−1^ following far-UVC subjection at 25% of the TLV for safe human exposure to 222 nm within a classroom scenario. This far-UVC driven SARS-CoV-2 aerosol inactivation dose-response relationship demonstrates the ability of current far-UVC technology, implemented with sufficient excimer lamps, to achieve substantial infectious airborne viral inactivation within occupied room environments and provides a foundation to design and implement far-UVC devices in buildings.

### Supporting Information

- · Presents additional methodological protocols, expansion of select results and statistical significance, and supplementary figure/table: protocols for Vero E6 cell culture, SARS-CoV-2 propagation and plaque assay infectious SARS-CoV-2 quantification; justification of the use of a 2:1 ratio of human saliva and DMEM as the aerosolization media; calculation for measured irradiance scaling relative to modeled average irradiance and fluence rate; further explanation regarding the statistical significance of total and infectious viral decay under far-UVC off and far-UVC on conditions relative to far-UVC mediated viral decay, including genome damage; the far-UVC dose required to facilitate 2-log decay in infectious SARS-CoV-2 based on average irradiance; representative size-resolved particle size distribution of aerosols within the chamber (Figure S1); average and standard error of experimental RH values with statistical significance (Table S1) (PDF)

## Acknowledgements.

We thank Dr. Holger Claus and Vivian Belenky for their technical expertise regarding UV light measurement and modeling. We also thank Dr. Irfan Ullah for providing the initial SARS-CoV-2 Beta variant stock in mice lung homogenate. Additionally, the authors would like to express appreciation to Benjamin Fontes, Melissa James, and entire the Yale University Biosafety Team for coordinating approval of controlled aerosolization experimentation with a BLS-3 infection agent, as well as Vincent Bernardo and the Yale University Gibbs Machine Shop for the construction of the aerosolization chamber. Finally, we express our gratitude to Taylor Burke for his assistance in measuring far-UVC chamber irradiance in the BSL-3 lab setting.

