## Supplemental Information for "The Susceptibility of Airborne SARS-CoV-2 to Far-UVC Irradiation"

**Supplementary Information**

Darryl M. Angel^1‡^, Irvan Luhung^1‡^, Keyla S. G. de Sá^2^, Jordan Peccia^1*^

^‡^ Equal Contribution

1 Department of Chemical and Environmental Engineering, Yale School of Engineering and Applied Science, Yale University, New Haven, CT, USA

2 Department of Immunobiology, Yale School of Medicine, New Haven, CT, USA

**MATERIALS AND METHODS**

**Viral Propagation and Quantification.**

*Cell Culture.*

Within a BSL-2 laboratory, Dulbecco’s Modified Eagle Medium (DMEM) (Gibco, #11965092) supplemented with 10% heat-inactivated fetal bovine serum and 1mM sodium pyruvate was prepared.(1) Vero E6-ACE2-TMPRSS2 cells were cultured in 75 cm^2^ culture flasks, with 10 mL of supplemented DMEM and 10 µL of 10 mg mL^-1^ puromycin, at 37°C in a 5% CO_2_ atmosphere.(1, 2) Vero E6 cells were passaged twice a week at a sub-cultivation ratio of 1:5, for flask re-seeding or well plate seeding, as instructed in Jureka and Basler (2002).(1) Three days before plaque quantification of infectious SARS-CoV-2, 2.0 x 10^5^ live cells were suspended in 2 mL of supplemented DMEM and plated into individual wells within 6-well plate.

*SARS-CoV-2 Propagation and Enumeration.*

Working stocks of SARS-CoV-2 Beta variant (B.1.351), originally obtained from infected K18 mice lung homogenate, were propagated using Vero E6 cells as outlined in previous methods.(1, 3) Second passage viral stocks, stored in non-supplemented DMEM at ~10^8^ PFU/mL, were used for experimentation.

For enumeration following chamber aerosolization experimentation, infectious and total SARS-CoV-2, collected in SKC BioSamplers, were quantified via plaque assay and ddPCR methods, respectively. Plaque assay infectious viral quantification was conducted using the Microcrystalline Cellulose (MCC) overlay method.(1) Serial dilutions of viral samples (0.4 mL) were incubated on Vero E6 cells in a 6-well plate for one hour, gently rocking every 15 minutes. Two mL of 2.4% MCC and supplemented DMEM (1:1 ratio) were subsequently added to the

well-plates and incubated for another 72 hours. All incubation steps occurred at 37°C in a 5% CO_2_ atmosphere. Following the 72-hour incubation, cells were fixed to the plate surface using 4% paraformaldehyde (#15735-20S, Electron Microscopy Sciences) before staining with a 0.5% Crysal Violet solution to visualize plaques.(1)

For total viral enumeration, DNA/RNA shield (#R1200, Zymo Research) was added to viral impinger samples, in a 1:1 ratio, to preserve viral RNA and inactivate SARS-CoV-2 prior to removal from the BSL-3 lab. Samples were stored overnight at -80℃, extracted using the Quick-RNA Viral Kit (#R1035, Zymo Research), and quantified for total SARS-CoV-2 RNA using ddPCR methods. The SARS-CoV-2 RNA extraction and ddPCR protocols have been previously reported.(4)

**Aerosolization Media.**

There are indications that proteins or other organic components in saliva may influence airborne virus exposure to far-UVC, with a prior study showing that aerosolization in artificial saliva significantly reduced far-UVC susceptibility of the aerosolized enveloped Phi6 virus relative to a media of sodium chloride and magnesium sulfate buffer.(5) The opted aerosolization media of a 2:1 ratio (disinfected human saliva:SARS-CoV-2 in DMEM) is meant for a closer representation of human generated viral aerosols in saliva relative to pure DMEM.

**Measured Irradiance Scaling.**

Across the vertical planes measured within the chamber, the irradiance at the center points in the closest (7 cm), middle (21 cm), and furthest (33 cm) measured locations from the excimer lamp are 7.69, 1.26, and 0.55 μW cm^-2^, respectively. Utilizing the quotient of the irradiance values for the center point of each of the three vertical planes from UV simulation

software (no semi-transparent plastic sheet) and the reported irradiance of the center points obtained from direct light meter measurements (includes semi-transparent plastic sheet), plane-specific scaling factors of 4.56 x 10^-2^, 4.11 x 10^-2^, 4.23 x 10^-2^ were calculated for chamber irradiance at 7, 21, and 33 cm from the far-UVC light source, respectively. These scaling factors were applied to each of the modeled vertical panels to obtain a higher resolution of the spatial distribution of irradiance within the chamber, accounting for the light dampening effect due to the semi-transparent plastic sheet. The scaling factor for the overall center point within the chamber (4.11 x 10^-2^) was further used to scale the simulated average irradiance and fluence rate within the chamber.

**RESULTS.**

**Relative Humidity.**

There is no statistically significant difference within the two experimental RH conditions when comparing average RH for far-UVC off and far-UVC on conditions (p > 0.05). However, there is a significant difference in the average relative humidities across the low and high RH conditions, confirming that the two experimental RH scenarios were sufficiently different (*p* < 0.01) (Table S1).

**SARS-CoV-2 Susceptibility to Far-UVC.**

*Experimental Total and Infectious Viral Decay Rate Constants.*

The total viral aerosol decay rate constants (± standard error) for far-UVC off and far-UVC on conditions are 15.9 ± 2.1 hr^−1^ and 20.8 ± 4.5 hr^−1^, respectively, at low RH and are 18.7 ± 2.1 hr^−1^ and 22.9 ± 3.1 hr^−1^, respectively, at high RH. Within experimental RH conditions, there is not a statistically significant difference in the total decay rates from background to far-UVC on

conditions (*p* = 0.32 under low RH; *p* = 0.28 under high RH). This result demonstrates that far-UVC did not contribute to viral aerosol deposition in the chamber. Furthermore, this lack of statistical significance based on ddPCR data suggests that additional factors which may contribute to ddPCR quantified SARS-CoV-2 loss, such as RNA damage specifically within the N1 region quantified, are insignificant at the far-UVC doses considered relative to the total loss rate of airborne viruses due to deposition within the experimental chamber. In contrast, the experimentally derived first-order infectious SARS-CoV-2 aerosol decay rate constants at low RH are 23.0 ± 2.6 hr^−1^ and 43.3 ± 2.7 hr^−1^ under far-UVC off and far-UVC on conditions (*p* ≤ 0.01), respectively, indicating that far-UVC technology facilitates the inactivation of infectious SARS-CoV-2 aerosols. There was also an increase in the infectious SARS-CoV-2 aerosol decay rate constants, from 24.1 ± 2.3 hr^−1^ to 51.6 ± 3.5 hr^−1^, under background to far-UVC on conditions at high RH (*p* ≤ 0.01).

*Far-UVC Inactivation as a function of Dose.*

In the absence of time-matched bioaerosol samples, the far-UVC facilitated log inactivation and viral aerosol surviving fractions can be calculated based on UV disinfection modeling.(6) Using first-order kinetics for single-stage decay, the fractional survival can be calculated from the far-UVC dose, determined by multiplying the average fluence rate or irradiance and exposure time, and the susceptibility constant (accurate up to a 2-log decay).(6-9) Within the experimental system utilized in this study, the far-UVC dose required to facilitate a 2-log (99%) reduction of exposed SARS-CoV-2 aerosols is 1.04 mJ cm^-2^ under the low RH scenario and 0.68 mJ cm^-2^ under the high RH scenario (0.76 and 0.50 mJ cm^-2^ irradiance-based doses, respectively).

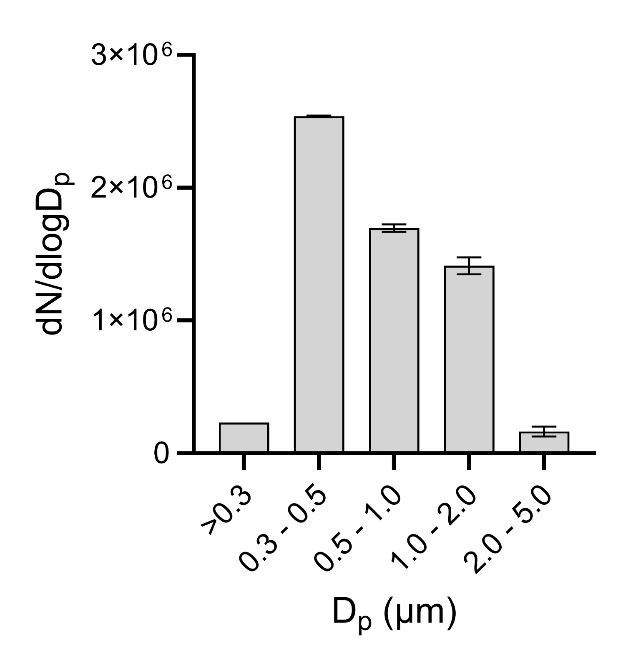

**Figure S1. Representative size-resolved particle size distribution of aerosols within the chamber.** Aerosols were generated by the atomizing nozzle during initial optical particle counter sampling averaged across low and high RH conditions (± standard error). Aerosols generated from 1:2 ratio of SARS-CoV-2 in DMEM and human saliva respectively

**Table S1.** Average and standard error of experimental RH values utilizing Kruskal Wallis followed by Dunn's Multiple Comparison test to assess the statistical significance between RH levels under far-UVC off and far-UVC on conditions.

| RH | Background Average (%) | Background SE (%) | Far-UVC On  Average (%) | Far-UVC On  SE (%) | P-Value (Background v. Far-UVC On) |
| --- | --- | --- | --- | --- | --- |
| Low | 38.0 | 0.3 | 39.2 | 0.2 | *p* > 0.05 |
| High | 66.7 | 0.5 | 67.5 | 0.4 | *p* > 0.05 |

**REFERENCES.**

1.	Jureka, A. S.; Basler, C. F. Propagation and quantification of SARS-CoV-2. *Methods Mol Biol* **2022,** *2452*, 111-129.

2.	Kinoshita, H.; Yamamoto, T.; Kuroda, Y.; Inoue, Y.; Miyazaki, K.; Ohmagari, N.; Tokita, D.; Nguyen, P. H. A.; Yamada, S.; Harada, S.; Kanno, T.; Takahashi, K.; Saito, M.; Shirato, K.; Takayama, I.; Watanabe, S.; Saito, T.; Ebihara, H.; Suzuki, T.; Maeda, K.; Fukushi, S. Improved efficacy of SARS-CoV-2 isolation from COVID-19 clinical specimens using VeroE6 cells overexpressing TMPRSS2 and human ACE2. *Sci Rep-Uk* **2024,** *14* (1).

3.	Rodrigues, T. S.; de Sa, K. S. G.; Ishimoto, A. Y.; Becerra, A.; Oliveira, S.; Almeida, L.; Goncalves, A. V.; Perucello, D. B.; Andrade, W. A.; Castro, R.; Veras, F. P.; Toller-Kawahisa, J. E.; Nascimento, D. C.; de Lima, M. H. F.; Silva, C. M. S.; Caetite, D. B.; Martins, R. B.; Castro, I. A.; Pontelli, M. C.; de Barros, F. C.; do Amaral, N. B.; Giannini, M. C.; Bonjorno, L. P.; Lopes, M. I. F.; Santana, R. C.; Vilar, F. C.; Auxiliadora-Martins, M.; Luppino-Assad, R.; de Almeida, S. C. L.; de Oliveira, F. R.; Batah, S. S.; Siyuan, L.; Benatti, M. N.; Cunha, T. M.; Alves-Filho, J. C.; Cunha, F. Q.; Cunha, L. D.; Frantz, F. G.; Kohlsdorf, T.; Fabro, A. T.; Arruda, E.; de Oliveira, R. D. R.; Louzada-Junior, P.; Zamboni, D. S. Inflammasomes are activated in response to SARS-CoV-2 infection and are associated with COVID-19 severity in patients. *J Exp Med* **2021,** *218* (3).

4.	Angel, D. M.; Gao, D.; DeLay, K.; Lin, E. Z.; Eldred, J.; Arnold, W.; Santiago, R.; Redlich, C.; Martinello, R. A.; Sherman, J. D.; Peccia, J.; Godri Pollitt, K. J. Development and application of polydimethylsiloxane-based passive air samper to assess personal exposure to SARS-CoV-2. *Environ Sci Technol Lett* **2022,** *9* (2), 153-159.

5.	Monika, D. E. A., Kondabagil K., Kunwar A. Comparative study of inactivation efficacy of far-UVC (222 nm) and germicidal UVC (254 nm) radiation against virus-laden aerosols of artificial human saliva. *Photochemistry and Photobiology* **2025**.

6.	Kowalski, W., *Ultraviolet Germicidal Irradiation Handbook: UVGI for Air and Surface Disinfection*. Springer: New York, 2009; p 1-501.

7.	Welch, D.; Buonanno, M.; Grilj, V.; Shuryak, I.; Crickmore, C.; Bigelow, A. W.; Randers-Pehrson, G.; Johnson, G. W.; Brenner, D. J. Far-UVC light: A new tool to control the spread of airborne-mediated microbial diseases. *Sci Rep-Uk* **2018,** *8*.

8.	Welch, D.; Buonanno, M.; Buchan, A. G.; Yang, L.; Atkinson, K. D.; Shuryak, I.; Brenner, D. J. Inactivation rates for airborne human coronavirus by low doses of 222 nm far-UVC radiation. *Viruses* **2022,** *14* (4).

9.	Buonanno, M.; Welch, D.; Shuryak, I.; Brenner, D. J. Far-UVC light (222 nm) efficiently and safely inactivates airborne human coronaviruses. *Sci Rep-Uk* **2020,** *10* (1).
